# Rapid divergence in vegetative morphology of a wind-pollinated plant between populations at contrasting densities

**DOI:** 10.1101/2021.07.09.451799

**Authors:** Jeanne Tonnabel, Patrice David, John Pannell

## Abstract

Plant sexual dimorphism is thought to evolve in response to sex-specific selection associated with competition for access to mates or resources, both of which will often be density-dependent. In wind-pollinated plants in particular, vegetative traits can have an important influence on both resource acquisition and the pollen dispersal and receipt, with potential conflict between these two components of fitness. We evaluated the role of plant density in shaping plant traits by measuring evolutionary responses in experimental populations of the sexually dimorphic wind-pollinated plant *Mercurialis annua*. After three generations of evolution, we observed divergence between high- and low-density populations in several vegetative traits, whereas there was no divergence for reproductive traits. A reversal in the direction of sexually dimorphic traits expressed in young plants evolved in both low- and high-density populations compared to the original population (stored as seeds). Compared to the source population, males at high density evolved to be taller when young, whereas at low density young females tended to become smaller. These results demonstrate that a simple change in plant density can induce rapid, age-dependent and sex-specific evolution in the ontogeny of vegetative organs, and illustrates the power of experimental evolution for investigating plant trait evolution.

## Introduction

Males and females of dioecious plants often differ in their morphological, life-history and physiological traits (Geber *et al*., 1999). Although sexual dimorphism in plants is rarely as extreme as that displayed by many animals (Lloyd & Webb, 1977), it has nevertheless evolved multiple times during angiosperm diversification. The South African genus *Leucadendron* provides a striking example of this convergence, with divergence in the degree of sexual dimorphism having evolved several times independently among species (Tonnabel *et al*., 2014). In other species, it has been found that sexual dimorphism may differ significantly among populations (e.g., *Silene latifolia*, Delph *et al*., 2002; *Rumex hastatulus*, Puixeu *et al*., 2019), suggesting evolutionary divergence in response to spatial and temporal variation in selection experienced by the sexes. A key factor that may vary dramatically among populations, and from one generation to the next, is plant density. Density is particularly interesting in the context of sexual dimorphism because we can expect it to modulate the strength of competition both for mates and for resources.

Although the role of sexual selection in shaping plant evolution was hotly debated for some time (Grant, 1995; Stanton, 1994), it is now well established that sexual dimorphism in plants may indeed be the outcome of competition for mates (Moore & Pannell, 2011; Lankinen & Karlsson Green, 2015). Following Darwin’s (1871) seminal work on the evolution of sexual dimorphism in animals, two modes of sexual selection that might bring it about have persisted in our thinking: selection resulting from male-male competition for access to females (‘intrasexual selection’); and selection resulting from the choice by females of their mating partners (‘intersexual selection’ or ‘female choice’). The possibility of a form of female choice in plants is theoretically possible but remains poorly evaluated (Tonnabel *et al*., 2021). In contrast, the role of intrasexual competition among males or hermaphrodites to fertilize ovules is well established (Moore & Pannell, 2011; Lankinen & Karlsson Green, 2015). Indeed, there is now a substantial body of theory exploring this notion (Arnold, 1994; Stanton, 1994; Tonnabel *et al*., 2019a), which is also supported by studies demonstrating the importance of male-male competition for trait evolution in plants (Bond & Maze, 1999; Delph & Herlihy, 2012; Schiestl & Johnson, 2013; Coccuci *et al*., 2014; Dorken & Perry, 2017; Lankinen *et al*., 2017), and, in dioecious species in particular, for trait divergence between males and females (Tonnabel *et al*., 2019a,b). Much of this work is consistent with Bateman’s third principle, which posits that male reproductive success is more limited by the number of mates they have access to, and therefore by intrasexual competition for mates, than female reproductive success (Bateman, 1948; Arnold, 1994; Stanton, 1994). Bateman’s third principle should apply more to populations in which many males compete with one another to pollinate a limited pool of females than in populations in which such competition takes place among fewer. As such, the importance of Bateman’s third principle should be sensitive to plant density, particularly in wind-pollinated plants in which mating depends on the spatial proximity of mates and other individuals competing for them (Tonnabel *et al*., 2019a,b).

While competition for mates can give rise to divergence between males and females for reproductive traits, vegetative traits can have a direct impact on the outcome of mating particularly in wind-pollinated plants in which both height and above-ground plant architecture can affect pollen dispersal and receipt (Klinkhamer *et al*., 1997; Eppley & Pannell, 2007; Pickup & Barrett, 2012; Harder & Prusinkiewick, 2013; Tonnabel *et al*., 2019a,b). For example, variation in plant height, branching patterns, branch length, and canopy diameter may affect the release and dispersal of pollen grains (Klinkhamer *et al*., 1997; Harder & Prusinkiewick, 2013). In the wind-pollinated herb *Mercurialis annua*, either elongated inflorescences or longer branches have been found to promote pollen dispersal over greater distances, increasing the number of a male’s mates (Eppley & Pannell, 2007; Tonnabel *et al*., 2019a,b). In *Leucadendron*, evolutionary transitions from insect to wind pollination are strongly associated with strong sexual dimorphism in vegetative traits (Tonnabel *et al*., 2014; Welsford *et al*., 2016), perhaps as a result of selection on plant architecture, or because males and females are no longer constrained to be recognized similarly by pollinators (Vamosi & Otto, 2002). As for reproductive traits, we should expect plant density to modulate the extent to which vegetative traits affect patterns of mating and thus the intensity of sexual selection.

Vegetative traits should not only have direct effects on mating success by influencing pollen dispersal, but they are of course of primary importance in the acquisition of resources. On the one hand, numerous studies have shown that the two sexes display different reaction norms to resource availability (e.g. water or nutrients) by allocating resources to their organs differently (reviewed in Tonnabel *et al*., 2017), suggesting that differing costs of reproduction between males and females may translate into sex-specific selection for accessing different resource components (Antos & Allen, 1990; McDowell *et al*., 2000; Harris & Pannell, 2008; Van Drunen & Dorken, 2012). Differences between the sexes in their resource needs are likely to be especially important in wind-pollinated plants, in which males produce large quantities of nitrogen-rich pollen (Harris & Pannell, 2008; Wright & Dorken, 2014; van Drunen & Dorken, 2012), whereas the production by females of seeds and fruits typically draw heavily on photosynthates and water (Antos & Allen, 1990; McDowell *et al*., 2000; van Drunen & Dorken, 2012). On the other hand, we should nevertheless expect vegetative divergence between the sexes to be limited by a common need to avoid losing the competitive race with neighbors to maintain adequate access to light: losing the race to harvest light will have deleterious effects on multiple plant functions (Labouche & Pannell, 2016). This limitation should apply less at low density, but at high density both males and females will typically need to adopt a similar architectural strategy (Labouche & Pannell, 2016; Tonnabel *et al*., 2017). Plastic responses to increased competition for light widely consist of a ‘shade avoidance’ strategy in both sexes, including increased plant height and inter-node elongation (Schmitt & Wulff, 1993; Sleeman *et al*., 2002). Thus, although males and females might differ in important ways in their needs for different resources, the extent to which they can afford do diverge will depend on the intensity of competition for light with neighbors.

The effect of density on the different components of plant fitness is evidently complex, but it seems clear that the direction of selection on reproductive and particularly vegetative traits is likely to differ between low- and high-density situations. Densities experienced by plant populations often vary over orders of magnitude in space and time (e.g. Boe *et al*., 2019), so that we should naturally expect to observe both plastic changes in morphology as well as evolutionary changes in morphological traits in response to exposure to different densities. Here, we explore the evolutionary responses to varying plant density in the wind-pollinated dioecious annual herb *Mercurialis annua* using experimental evolution. We established five independent replicate populations at high density and five at low density, and allowed each population to evolve under the influence of natural and sexual selection over the course of three generations. We manipulated plant density by varying the density of pairs of males and females. Our experimental populations were established using the seeds from a source (experimental) population with high genetic variance for several plant traits of interest (see below). We thus followed classical procedures of experimental evolution, whereby potentially divergent selection is allowed to act under contrasting environmental conditions on the standing genetic variation sampled from a common founding population (Kawecki *et al*., 2012). While the chief focus in such studies is typically the comparison between diverging lines after a given time or number of generations of evolution, in some organisms it is possible to compare evolved populations to a source population maintained in a dormant state (e.g., by freezing, which is possible for microbes; Woods *et al*., 2011). We were able to draw such a comparison by maintaining a stock of seeds from the source population in a fridge, allowing us to assess the direction in the observed evolutionary responses under both density treatments.

We designed our experiment based on previous work on different populations of *M. annua* that suggested that plant spacing in our low-density treatment should bring about a largely monogamous mating system whereas mating at high density would involve intense competition among males for siring success (Eppley & Pannell, 2007). In the first generation of our experiment, we conducted paternity analyses on two populations with contrasting density and confirmed substantial differences between them in mating patterns (Tonnabel *et al*., 2019a,b). Specifically, females in the low-density treatment were substantially more likely to mate with their paired male than those in the high-density treatment; however, even at low density approximately 60% of seeds were still sired by extra-pair males – i.e. the mating system was still far from monogamous (Tonnabel *et al*., 2019b). We initially expected that the high-density treatment should intensify competition among males for fertilizing ovules on both their paired female as well as on other females nearby, so that intra-sexual selection should select for large males that produce more pollen dispersed over a greater number of female mates; in contrast, we expected that the low-density treatment would cause females to mate primarily with their paired male (i.e., mating would be largely monogamous), so that selection should favor males that ceded resources to their paired female (e.g., by growing small), thereby allowing her to produce more ovules to which the paired male would have privileged access. Male-female pairing is known to be associated with dwarf males in a number of taxa, both animals (Vollrath & Parker, 1992; Coddington *et al*., 1997; Vollrath, 1998; Kuntner & Elgar, 2014; Rouse *et al*., 2015) and plants (mosses – Hedenäs & Bisang, 2011). As will be seen, and consistent with a reduction, instead of the initially intended suppression, of extra-pair matings at low density, our results were more subtle. We observed significant responses to selection over the three generations of evolution, some of which differed from our initial expectations, illustrating the complexity of selection when traits affect both competition for resources and for mates.

## Materials and methods

### Study system and seed origin

*Mercurialis annua* is an annual wind-pollinated herb distributed throughout southern and central Europe and around the Mediterranean Basin (Tutin *et al*., 1964). The species complex includes dioecious, androdioecious and monoecious populations located in different parts of its range (Durand, 1963; Pannell *et al*., 2004). For the current experiment, we focused on dioecious populations that naturally exhibit strong sexual dimorphism, with males being shorter than females and displaying stalk-like (pedunculate) inflorescences that enhance pollen dispersal (Harris & Pannell, 2008; Tonnabel *et al*., 2019b). Both sexes start producing flowers shortly after seed germination. Growth is indeterminate and reproduction continues until environmental conditions deteriorate, and plants die (Pannell, 1997a).

We began our experiment by pooling seeds from 35 populations of *M. annua* sampled from northern Spain, with seed families sampled from approximately 30 females per population (see Tonnabel *et al*., 2017, 2019b). *M. annua* displays a metapopulation structure and dynamic characterized by frequent events of colonization and extinction (Obbard *et al*., 2006; Eppley & Pannell, 2007); our experimental source population therefore represents the genetic and phenotypic variation present in wild populations at the metapopulation level. Before our experiment, we grew plants from the pooled source population in a common garden in Lausanne for three generations under uniform growing conditions (from 2012 to 2014) at an intermediate density compared to our later density treatments. This common garden was aimed at reducing any maternal effects and/or genetic correlations caused by population subdivision across the metapopulation. We refer to seeds harvested after these three generations in the common garden as the source population. These seeds from the source population served two purposes in our experiment: (1) setting up our ten experimental populations with individuals drawn from a common pool and thus with a similar genetic composition; and (2) comparing the evolved traits in the low- and high-density conditions to the source population under common environmental conditions in a common garden to inform on the direction of evolutionary changes. Seeds from the source population were stored in a fridge at 23°C with a humidity of 58% for ensuring their conservation during the three years of the experiment.

### Experimental evolution protocol

We established ten experimental populations of *M. annua* divided into five populations that evolved independently from one another at low density and five others at a high density in semi-natural conditions at the experimental field platform of the LabEx CeMEB in Montpellier, France. Each of the ten experimental populations were composed of 100 males and 100 females and were maintained at their assigned density for three generations (grown in spring of 2015, 2016 and 2017; see Tonnabel *et al*., 2017, 2019a,b for a description of the first generation), giving a total of 2,000 plants grown each generation. Each garden consisted of a square array of 10 × 10 pots, each containing one male and one female growing together, and therefore competing for light (and other nutritive resources). Each of our experimental populations was established using seeds from the source population, following the classical approach adopted in studies of experimental evolution (Kawecki *et al*., 2012). We allowed plants to mate naturally in each of their populations (see below), then collected all seeds at the time of harvest, bulked the seeds of all females (within each population separately), and subsequently sowed seeds for the following generation by randomly sampling individuals from the bulked sample. This procedure ensured that each plant contributed, on average, to the following generation proportionally to the number of seeds it produced or sired.

At the beginning of each generation, seeds were individually germinated in greenhouses using pots in sterile compost. In the first generation, all seedlings came from our source population; in this first generation, seedlings were assigned to one of the ten experimental populations and these populations were kept separate from that point on. Prior to being transplanted into their populations in the field sites, seedlings were distributed randomly in space across the greenhouses and their positions were shuffled frequently. Seedlings were grown until most plants had reached maturity and could be sexed. At this point, after approximately 1.5 months of growth, pairs of males and females of the same experimental population were transplanted into 2L pots of 20cm of diameter containing sterile soil (1/3 of sieved clay and chalky soil, 1/3 of recycled compost and 1/3 of compost). These male-female pairs were moved outside and assigned to their respective experimental population. During the first generation, males and females were paired accordingly to their size to decrease asymmetries in competition caused by differences in the timing of germination (Tonnabel *et al*., 2017). To increase the possibility of strong asymmetrical competition for light within pairs, we slightly changed the protocol in the second and third generations in which we paired males and females randomly.

Because our initial hypotheses were focused on the effects of male-male competition and intra-pair competition for light, we attempted to minimize differences in competition for light among pots from the two different density treatments. We thus allowed all plants to grow under the same growing conditions for most of their growing period. Therefore, we imposed different densities only for a brief period of mating prior to harvest. Specifically, all experimental populations were first grown at a low-density corresponding to a pot separation of 1.0 m. Approximately a month after transplanting the plants outside (with slight variation due to between-year variation in weather conditions), we changed the position of all plants in the experimental populations while maintaining between-pot distances of 1.0 m and 20 cm for low- and high-density respectively (between-pot distances here and below designate distances between pot centers). Plants were allowed to continue to mate in their new positions for four weeks. After these four weeks, and for each population separately, we harvested all female plants and bulked them in drying bags. After drying all females separately for each population, we separated vegetative parts from seeds, which were kept to establish the next generation. As fruits disperse their seeds several days after fertilization, all seeds harvested after four weeks should have been fertilized during the phase of differential density application. Low adult mortality was observed in each generation.

Our experimental field consisted of two rows of five sites each. At each generation, we randomly assigned experimental populations among these ten sites by following two rules: we assigned (1) either two or three replicates of each treatment to each of the two rows and (2) each column contained one replicate of each treatment. This block design aimed at minimizing any differential effect of possible environmental gradients on populations belonging to the two different treatments. The 10 m by 10 m sites were separated by 20 meters in order to reduce gene flow between populations. Potential pollen flow was likely further reduced by the growth of dense vegetation in the meadow between sites (the growth of vegetation between pots within populations was prevented by a tarpaulin). Previous work on *M. annua* suggests that most mating occurs over short distances (Eppley & Pannell, 2007; Hesse & Pannell, 2011). We confirmed this small-scale spatial pattern of mating on the basis of pollen dispersal kernels estimated for two of our experimental populations (Tonnabel *et al*., 2019b).

### Assessment of evolutionary responses to selection

After three generations of evolution, individuals from all experimental populations, plus those from the original source population (maintained in a fridge), were established in a single common garden at the experimental field platform of the LabEx CeMEB in Montpellier, France. Seeds were initially germinated in greenhouses by adopting the growing procedures described above. When plants had reached maturity, they were transplanted to the common garden in individual pots. The garden consisted of two blocks, one of 21 × 21 plants and the other of 20 × 22 plants, with an extra plant placed at one corner, giving a total of 441 plants per block. In both blocks, plants were grown in 2 L pots of 20 cm of diameter, placed with a between-pot distance of 40 cm (a density that was intermediate between our two experimental densities). Some mortality occurred in the course of growth in the garden, which led to a dataset including 708 plants. Although this mortality prevented us from obtaining a fully balanced block design at the end of the experiment, all sexes and experimental populations were represented in each block, except for one population for which a manipulation error led to solely males being placed in both blocks. We recorded the coordinates of each plant within the two experimental blocks.

To assess male and female vegetative growth, we recorded plant height as the distance between the soil and the highest pair of leaves: (1) at the time of transplantation, (2) three weeks after transplantation, and (3) at the time of the final harvest (see below). In the following, we refer to our three repeated measures of plant height as ‘young’, ‘intermediate’, and ‘old’. For these plant height measurements, we excluded the length of exert male peduncles to maintain comparability between the sexes. Three months after germination, all plants were harvested and measured. For all plants, we recorded its sex, the diameter of its canopy (as the longest horizontal length found between two leaves), and the length of the first two branches (i.e., the lowest ramifications down the plant whose length we averaged and which value we refer to as branch length in the following), and its above-ground vegetative biomass.

For males, we recorded the total number of pedunculate inflorescences, as well as their biomass, which we used as a proxy for pollen production (pollen accounts for 60% of male flower biomass; Pannell, 1997b). On fresh male plants, we also measured the length of the five inflorescence-bearing peduncles sampled on the fifth highest nodes of the primary axis. These five measures were later averaged, yielding our estimate of peduncle length. For females, we separated vegetative and reproductive tissues after drying the whole plant. We further weighed the seed mass and used an automatic seed counter (Elmor C3; Elmor Angewandte Elektronik, Schwyz, Switzerland) to assess seed number and size.

### Statistical analysis

To test for sex-specific evolution in plant architecture, we constructed full linear mixed-effects models (LMM) including one focal morphological trait as the response variable and sex and treatment (comprising the source population, the high- and low-density populations, as explained below) and their interaction as independent variables. All following analyses were performed separately for each morphological trait. We accounted for other sources of variance in our LMMs by including, as random effects: (1) our two experimental blocks as implemented in the common garden; and (2) the experimental populations for which the random effect was treated as sex-specific. First, we used likelihood ratio tests (LRT) to establish differences in the strength and/or direction of sexual dimorphism between the evolved populations, and between the source and the evolved populations; here, we compared our full LMMs to those in which the sex by treatment interaction had been removed. This first test informs on the presence of sex-specific evolutionary responses. For variables in which such a sex-specific response was observed, we tested for trait differences between sexes by comparing models explaining the trait and including or not sex as a fixed variable for each treatment type separately using LRTs (including the same random structure as described above). Second, we tested whether the mean values of morphological traits had changed between the evolved populations, and between the source and the evolved populations, regardless of sex; here, we compared models explaining morphological traits that contained sex and treatment as response variables to nested models that excluded the treatment effect. This second test informs on the presence of an evolutionary response aligned between sexes. Only this second test was performed for reproductive traits (because such measures are sex-specific), with the slight difference that populations were treated as random effects in separate models for males and females.

To assess overall differences between evolved populations and between the source, as well as pairwise differences between the three different population types, we performed four different LRTs for each focal morphological trait: including in the treatment factor either the three categories (source, high-density and low-density) or their pairwise combinations (see Table 1). We applied the same statistical procedure to the ‘reproductive effort’, computed as the biomass of the reproductive parts (i.e., seeds or male inflorescences) divided by the plant’s vegetative biomass. Unless otherwise specified, we fitted all models using the R package ‘lme4’ version 4_1.1-21 (Bates *et al*., 2015) in R version 3.6.1 (R Core Team, 2019).

**Table 1:**
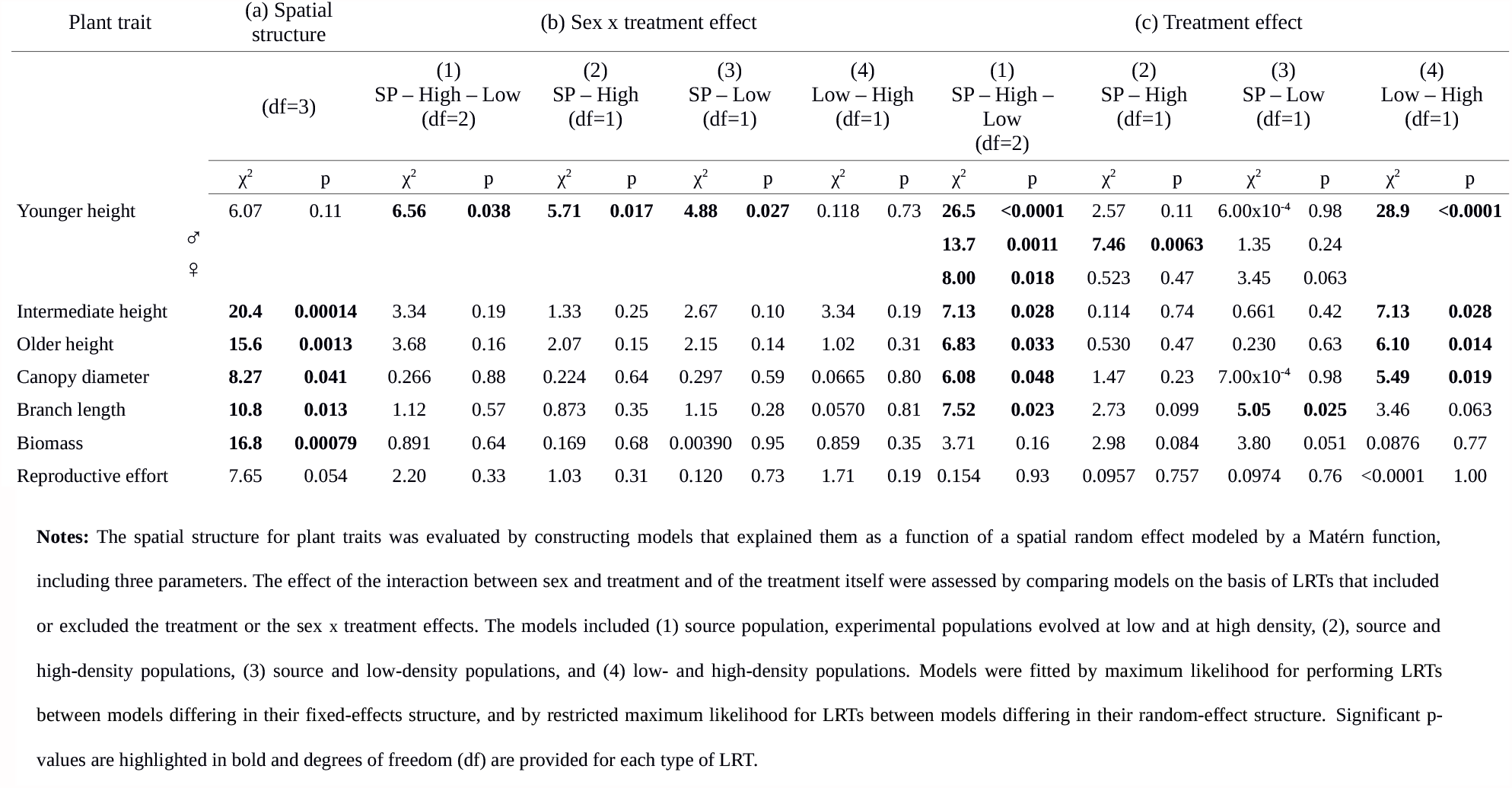
Testing for (a) spatial structure in plant vegetative traits, (b) sex-specific evolutionary response, and (c) non-sex-specific evolutionary response in these vegetative traits of *Mercurialis annua* plants that evolved at high- *versus* low-density and compared to our source population (SP) over the course of three generations, as assessed in a common garden.

Because environmental variation occurring within our semi-natural common garden arrays could have elicited spatial variation in both vegetative and reproductive traits, we included in our LMMs an additional random effect describing the spatial distribution of the trait within the common garden array, implemented as a Matérn correlation function, which models autocorrelation as a function of distance between plants, fitted using the R package ‘spaMM’ version 2.6.39 (see Rousset & Ferdy, 2014 for further details). We then used LRTs to compare models with or without the spatial random effect; this analysis revealed that several traits were indeed subject to a spatial structure (Table 1). For these traits, we repeated the statistical procedure described above, testing the effects of the treatment and treatment by sex interactions in models that included the effects of spatial structure. Because our results were robust to the inclusion of a spatial random factor explaining plant trait variation in LMMs for each of those variables that revealed significant spatial structure (see Table 1), we present only results of non-spatial LMMs as implemented in the R package ‘lme4’ (Bates *et al*., 2015). We compared the above statistical models either by maximum likelihood, or by restricted maximum likelihood when they differed in their fixed effects or in their random structure, respectively. Predicted values of vegetative traits for each sex and in each population types (source, low-density and high-density) are either shown in Figures 1 and S1, or in Table 2.

**Table 2:** Predicted sex-specific vegetative traits (intermediate height, canopy diameter, biomass and reproductive effort) in evolved and source populations (SP) of *Mercurialis annua* grown in a common garden after three generations of evolution.

|  | Intermediate height | Canopy<br>diameter | Biomass | Reproductive effort |
| --- | --- | --- | --- | --- |
| SP - Females | 29.7 (±0.935) | 11.5 (±0.681) | 5.09 (±0.303) | 0.0783 (±0.00969) |
| SP - Males | 25.9 (±0.776) | 9.71 (±0.541) | 3.52 (±0.269) | 0.170 (±0.00798) |
| Low - Females | 27.6 (±0.439) | 11.2 (±0.267) | 4.80 (±0.207) | 0.0831 (±0.00363) |
| Low -Males | 26.0 (±0.405) | 9.89 (±0.235) | 3.20 (±0.202) | 0.170 (±0.00315) |
| High - Females | 29.1 (±0.428) | 11.8 (±0.257) | 4.74 (±0.206) | 0.0884 (±0.00345) |
| High - Males | 26.7 (±0.413) | 10.4 (±0.244) | 3.29 (±0.203) | 0.166 (±0.00330) |

**Notes:** The null models predicted each response variable as a function of sex, treatment and their interaction and included both block and sex by population random effects.

**Figure 1.**
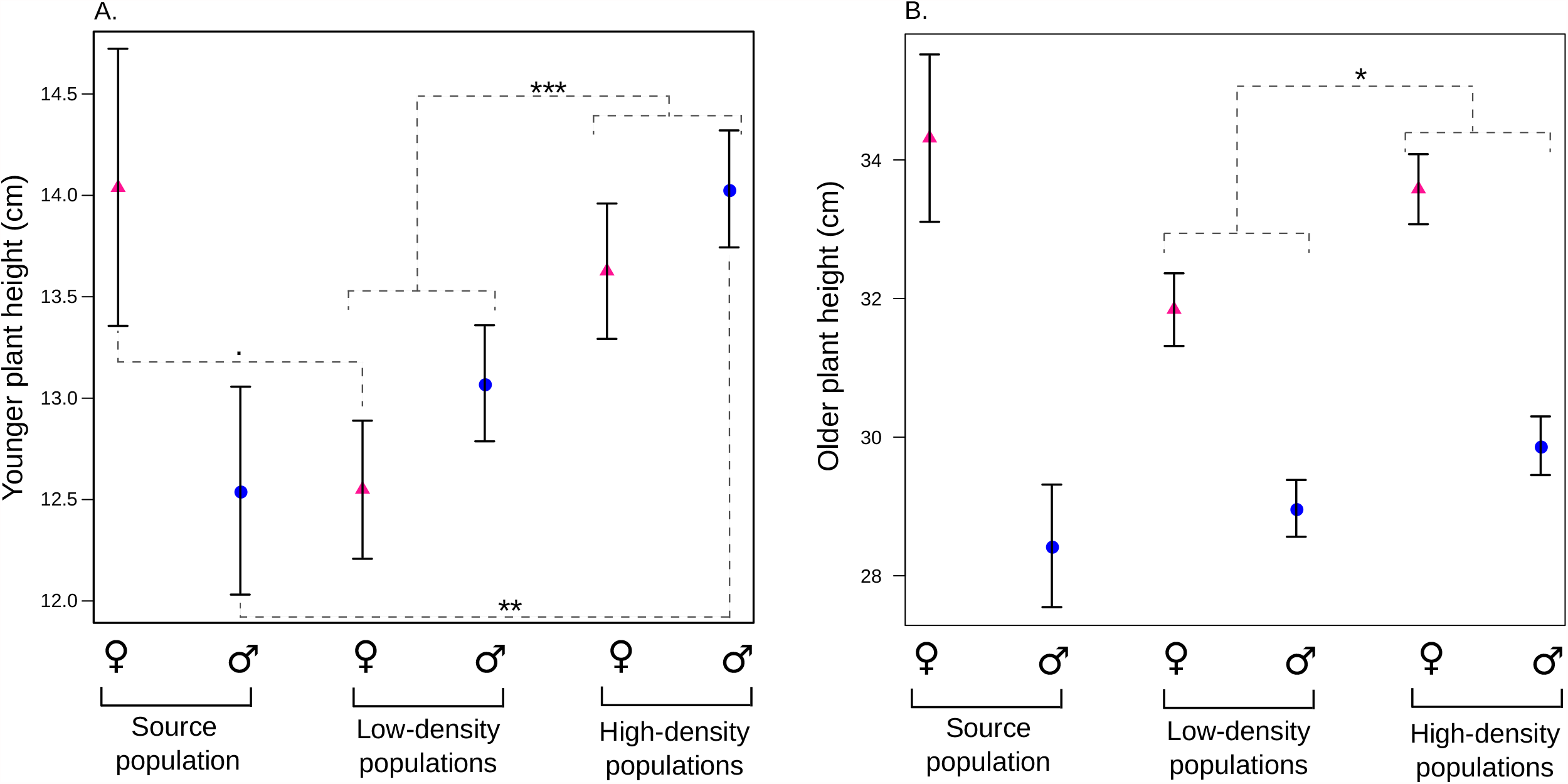
Predicted sex-specific plant height in evolved and source populations of *Mercurialis annua* grown in a common garden after three generations of evolution. (a) Younger plant height and (b) older plant height were treated as response variables in our null models, which included both block and sex by population random effects. Females and males are represented by pink triangles and blue circles, respectively. The significance of differences between treatments (source, low-density and high-density) in models combining both sexes and in sex-specific models was evaluated using LRTs (.*p* < 0.10, \*\**p* < 0.01, \*\*\**p* < 0.001). Separate models between sexes were performed only for the younger plant height (for which the sex by treatment interaction was significant). Horizontal bars indicate standard errors in model estimates.

To test whether our density treatment elicited differences in the sex-specific trait correlation structure, we first computed for each sex and treatment the Pearson correlation coefficients and associated p-values between all pairs of traits. We then applied a fourth-order tensor analysis, initially designed to compare *G* matrices (Hine *et al*., 2009; Aguirre *et al*., 2014), to compare the three matrices of phenotypic correlation obtained for each sex and corresponding to our three treatments (Hine *et al*., 2009; Aguirre *et al*., 2014). This tensor analysis allows simultaneous comparisons of the complete multidimensional variation between matrices while retaining uncertainty in correlation estimates (Hine *et al*., 2009; Aguirre *et al*., 2014). It estimates matrices and eigentensors, whose elements summarize independent aspects of how the original correlation matrices differ from one another. Each independent eigentensor accounts for a fraction of the between-matrix differences and is composed of eigenvectors corresponding to linear combinations of changes in pairwise correlations between traits. We used bootstraps to capture variation in the uncertainty in estimating phenotypic correlation matrices for the source, low-density and high-density populations separately; here, we performed 1,000 bootstraps for each treatment, each time sampling a different set of plants to estimate correlations while keeping the number of plants constant. We then compared the three phenotypic matrices obtained at each bootstrap iteration by performing a tensor analysis, as implemented in the package Gmatools version 0.37 (Chantepie, 2016). For each sex separately, we represented the mean position (and its credibility interval at 95%) of each treatment along eigentensors that explained >80% of the variation between the three compared matrices.

## Results

After three generations of evolution, plants evolved under high density diverged from those evolved under low density for several vegetative traits that involved greater allocation to growth (Table 1; Fig. 1b). These responses were common to both sexes (Table 1; Fig. 1b). In particular, both males and females that evolved at high density displayed greater plant height (at all ages) and greater canopy diameters than plants at low density (Table 1, 2; Fig. 1a,b). In contrast, there were no evolutionary divergence among treatments for plant biomass or for reproductive effort (Table 1), nor were there any treatment differences for reproductive traits measured separately on males and females (i.e., peduncle length, number of peduncles, number of peduncles above the plant, peduncle mass, seed number, seed size, total seed mass; Table S1).

Plants from the low- or high-density populations also diverged from the source population in two vegetative traits: young plant height and branch length (Table 1). The changes in young plant height differed between sexes to the extent of manifesting a reversal in the direction of dimorphism, as revealed by significant sex x treatment interactions (Table 1, 2; Fig. 1a). These interactions remained significant when the experimental populations of both or only of a single density treatment were included along with the source population (Table 1). When young, females were taller than males in the source population, as is typical for *M. annua* (e.g. Harris & Pannell, 2008), but the reverse pattern was found for the evolved populations of both the low- and high-density treatments (Fig. 1a). This reversal in the direction of sexual dimorphism can be attributed both to males from the high-density populations evolving to be ∼1.5 cm taller than those from the source population (note the significant effect of the source *versus* the high-density origin in models considering only males; Table 1; Fig. 1a), and to females from the low-density treatment evolving to be ∼1.5 cm shorter than those from the source population (note the marginally significant effect of the source *versus* low-density treatments in models that included only females; Table 1; Fig. 1a). These evolutionary changes resulted in significant differences between sexes in young plant sizes at both densities, as revealed by a significant effect of sex in models explaining young plant size for each density separately (low-density: *X*_i_^2^=7.62, df = 1, *P* = 0.006; high-density: *X*_i_^2^=12.0, df = 1, *P* = 0.001). Regardless of sex, plants that evolved at both densities also displayed shorter branches compared to the source population, but this effect was significant only for low-density populations, as revealed by a significant effect of the source *versus* low-density origin in models including both sexes (Table 1, 2; Fig. S1).

Our analysis revealed more similar trait correlations among the evolved lines than between them and the source population. Several morphological and/or reproductive traits were correlated with each other (significant pairwise Pearson correlation tests; Figure S2). Our tensor analysis confirmed a stronger and significant resemblance between correlation matrices of the two evolved lines than between them and the source population. In males, a single linear combination of trait correlations accounted for the vast majority of differences among the three correlation matrices (i.e., the first eigenvector of eigentensor 1 captured 81% of variation in eigentensor 1, which itself accounted for 91% of the variation among the three phenotype matrices). Males from both the low- and high-density populations differed in their position on this linear combination of traits compared to males from the source population (based on a comparison of 95% credibility intervals; Fig. S3a). This result suggests that the correlation structure was modified similarly in the evolved populations compared to the source population (Figure S2). In females, two linear combinations of trait correlations explained most differences among correlation matrices (i.e., the first two eigenvectors of eigentensor 1 captured respectively 53% and 42% of variation in eigentensor 1, which itself accounted for 99% of the variation among the three phenotype matrices). Along these two linear combinations of trait correlations, females from both low-density and high-density populations showed slightly divergent positions from the source population, albeit with overlapping credibility intervals (Fig. S3b).

## Discussion

Our experiment revealed rapid evolutionary responses to differences in plant density, with divergence in several vegetative traits between the two densities. After only three generations of divergent evolution, plants evolving at high-density differed from those evolving at low density, being taller at all ages and displaying a wider canopy in both sexes. Surprisingly, we also observed a reversal in the direction of sexual dimorphism in plant size in young plants in both evolved treatments compared to the source population. Our results show that sexual dimorphism in plant populations can evolve rapidly both in extent and direction, and illustrate the complex and unpredictable ways in which fitness through each of the two sexual functions maps to plant allocation and architectural phenotypes in the context of variation in density.

### The dual effects of density on competitive interactions

Our experiment was initially designed to test predictions concerning the effects of density on selection on plant traits via its modulation of the intensity of sexual selection among males for access to ovules. We predicted that higher density, by increasing the degree of polygamy compared to low density, would place more relative weight on male-male competition for mates. Accordingly, we estimated that the male opportunity for sexual selection (*i*.*e*., variance in the number of sexual partners) was elevated by 65% (0.43 *versus* 0.26) in one high-density population compared to another at low-density, while this metric remained unaffected by the change in density for females (0.11 *versus* 0.12), thereby suggesting that our high-density condition indeed exacerbated male-male competition (Tonnabel *et al*., 2019a). We reasoned that male-male competition for accessing ovules in populations at high density would ultimately favor increased pollen production and male morphologies that promote effective pollen dispersal. Of course, this reasoning in the context of our experiment rests on the assumption that mating patterns measured in the sub-sampled populations in the first generation (Tonnabel *et al*., 2019b) are representative of mating in the respective populations in generations 2 and 3, too. Although we could not check this assumption exhaustively throughout the experiment, the observed patterns of mating at high *versus* low density are consistent with expectations for the leptokurtic pollen dispersal kernels commonly estimated for wind-pollinated plants (Austerlitz *et al*., 2004; Goto *et al*., 2006; Gaüzere *et al*., 2013; Geber *et al*., 2014), including *M. annua* (Tonnabel *et al*., 2019b).

Notwithstanding the effect of density on mating opportunities, previous results on the first generation (Tonnabel *et al*., 2017) also pointed to higher competition for light at high than low density in both sexes. We should expect competition for light to change selection on females more than on males, if female reproduction relies more heavily on carbon. Accordingly, in the first generation, the female opportunity for overall selection (*i*.*e*., variance in the number of offspring) was increased by 50% (0.48 *versus* 0.32) in one high-density population compared to another at low-density, and this increment was not generated by competition for access to sexual partners (Tonnabel *et al*., 2019a). While it thus seems reasonable to suppose that the differences in density imposed by our experiment affected plant evolution through their effects both on patterns of mating and on competition for resources, our experimental design ultimately does not allow us to discriminate between these two likely effects.

### Possible responses to selection at different densities via effects on mate *versus* resource acquisition

From the arguments above, it seems reasonable to suppose that the differences in density imposed by our experiment affected plant evolution through their effects both on patterns of mating and on competition for resources. Below we discuss likely ways by which these two forms of competition may have contributed the evolutionary reponses we observed, keeping in mind that our design ultimately does not allow us to unambiguously attribute each response to one or the other (as we manipulated only density).

Some, but not all, of our results seem consistent with a response to selection on males for stronger access to mates. Plants mating at high-density evolved greater height and canopy diameter than those mating at low-density. Such responses are possibly the consequence of selection for mate acquisition by males, given that at least branch length has previously been shown to enhance pollen dispersal and mate acquisition in *M. annua* (Tonnabel *et al*., 2019b), and that plant height has long been considered important for promoting pollen dispersal in general (Klinkhamer *et al*., 1997; Harder & Prusinkiewick, 2013). The evolution of taller and larger plants at high density could alternatively be interpreted as a way to increase allocation to pollen production, as male flower number was strongly correlated with size traits (i.e., greater height, canopy diameter and branch length) in our populations. However, this interpretation is not consistent with the lack of divergence in reproductive effort and reproductive traits between plants growing at high *versus* low density. Thus, differential selection on mate acquisition is a more plausible explanation than selection on pollen production to explain the differences among treatments, as was previously predicted in this system by estimates of both sexual and fecundity selection (Tonnabel *et al*., 2019b).

Significantly, the evolution of larger canopy diameters and greater height for intermediate and older plants at high than at low density was not specific to males, i.e., there was no indication of a divergence in sexual dimorphism between the two densities. Such a parallel response to differences in density by the two sexes would be consistent with expectations under strong genetic correlations for the relevant traits between the sexes, e.g., with selection on males for more efficient pollen dispersal and a correlated response in females (a possible form of intra-locus genetic conflict between the sexes; Hosken *et al*., 2019). However, although sexual dimorphism at the juvenile stage did not diverge between high-density and low-density treatments in our experiment, it did change considerably compared to the source population (even to the extent of being reversed, albeit temporarily). This suggests that genetic correlations between the sexes are unlikely to have been much of a constraint on responses to selection in our experiment, at least at the juvenile stage, and they thus cannot easily explain the lack of difference in sexual dimorphism between the high- and low-density populations at older ages. Moreover, it seems that genetic correlations may themselves have changed over the course of our experiment, given that the structure of phenotypic correlations among traits were quite responsive to selection, having apparently evolved from the ancestral population in both groups of populations in our experiment. If we accept that inter-sexual genetic correlations did not pose a fundamental constraint on trait divergence between treatments in our experiment, it would seem that competition for light experienced by both sexes may have contributed to the observed parallel evolution of larger vegetative traits at high density.

### Evolution of a reversal in sexual size dimorphism between the two densities

Our experiment revealed a reversal in the direction of sexual dimorphism in plant size in young plants in both evolved treatments, compared to the source population. Given that our manipulations altered conditions experienced only later in life (recall that we imposed the density difference only after four weeks of growth), the expression of a response to selection by young plants may appear surprising. However, competitive ability late in life depends critically on resource allocation and physiological decisions taken much earlier, particularly because competition for light is strongly asymmetrical so that losing the competitive race early in life would have severe fitness implications later (Weiner, 1980). Nevertheless, it remains difficult to find a single explanation for a *reversal* in the direction of sexual size dimorphism in young plants at both densities, especially considering that the evolutionary trajectories leading to it, from a common initial state (represented by the source population), differed : predominantly, a decrease in young female plant height at low density, versus an increase in young male plant height at high density. One possibility is that this results from quite different modes of selection in the two density treatments. At low density, the evolution of shorter young females may reflect relaxed competition for light compared with the intermediate density experienced by the ancestral source population rather than differences in sexual selection, given that female reproductive success was independent of density, at least in the first generation of the experiment (Tonnabel *et al*., 2019b). High density, by contrast, intensified (rather than relaxed) competition for light in both sexes, and increased competition for mates specifically in males, to which populations may have responded by increasing growth in males (compared to the source population). Note that a male-specific increase in allocation to growth is not likely to result from competition for light only, given the weaker carbon needs of male than female reproduction in *M. annua* (Harris & Pannell, 2008).

Our explanation resonates to some extent with observations made for the wind-pollinated dioecious herb *Rumex hastatulus*. Pickup and Barrett (2012) and Puixeu *et al*. (2019) showed that males of *R. hastatulus* tend to be taller than females when pollen is dispersed, whereas females become the taller sex at the time of seed dispersal, consistent with a siring advantage of tall males. Similarly, in *M. annua*, the dispersal of pollen from inflorescences held above the plants or from longer branches has been shown to increase siring success (Eppley & Pannell, 2007; Santos del Blanco *et al*., 2019; Tonnabel *et al*., 2019b). While the timing of growth in height in *R. hastatulus* might seem to make more sense than our observations for *M. annua*, note that even a simple change in the competitive environment in *M. annua* could lead to age-dependent changes in the direction of sexual dimorphism. In our *M. annua* populations, females end up being taller than males at reproductive age even when they are smaller as juveniles. As is apparent in Fig. 1, the differences in relative sizes between the source population, the low-density evolved populations, and the high-density evolved populations for each sex at the juvenile stage were generally conserved at the reproductive stage, suggesting that plants can respond to selection applied during the reproductive period through changes in development at an earlier stage. To confirm the possibility that competition for mates late in a plant’s life affects early resource allocation, future experiments should address the link between age-dependent patterns of resource allocation and pollen dispersal (i.e., siring success).

### Correspondence between measured selection gradients and responses to selection

It is interesting that the evolutionary responses observed after three generations of selection under contrasting density treatments did not in general align with the selection gradients that we had measured in the first generation, in which selection seemed to favor taller, broader and heavier females at both densities (Tonnabel *et al*., 2019b). While we found a difference in these female traits between the two density treatments, female size did not in fact increase compared to the ancestral population (rather, a decrease was observed in low-density populations). Similarly, selection gradients in the first generation seemed to favor broader and longer-branched males at high density and longer peduncles at low density, albeit weakly, and male height was not favored at either density (Tonnabel *et al*., 2019b). Yet our results indicate that male height did in fact evolve (especially at high density and at the juvenile stage). Similar inconsistencies have been found in other studies that compared selection gradients and the results of experimental evolution, and have been attributed to patterns of standing genetic variation, heritability and pleiotropy (e.g. Gervasi & Schiestl, 2017). While strong heritabilities have typically been measured for plant height (e.g. Khan *et al*., 2018), artificial selection targeting this trait has also been found to drive changes in various other aspects of plant morphology, phenology and physiology (Zu & Schiestl, 2017). Unfortunately, we do not know the genetic variances and covariances for the traits we measured, and thus cannot speculate about such possibilities.

### Caveats and concluding remarks

Evolution of our experimental populations will have been a response to the divergent conditions under which they were placed in the two treatments, but also to general changes in the conditions experienced by all populations relative to those experienced by the ancestors of our source population prior to the experiment’s establishment. In particular, the fact that we harvested seeds at only one point in time and did not retain seeds for further generations that had been dispersed earlier likely gave rise to selection for late pollen and seed production, traits that may ultimately display genetic correlations with some of the traits we measured. Unmeasured maternal effects may also have impacted our results, not least because our evolved populations were cultivated for three generations in a different place and in different conditions than prevailed in the source population itself (though the latter had also grown in similar conditions for several generations). It is difficult to evaluate the respective impacts of adaptation to general experimental conditions and of maternal effects, but they may have contributed to the complexities in our results. Nevertheless, it seems unlikely that they acted in a sex- or treatment-specific manner. Finally, it is possible that differences in mating patterns between low- and high-density conditions, as documented by Tonnabel *et al*. (2019b), may have led to differences in the importance of genetic drift between them. Although treatment-specific genetic drift may ultimately compromise the interpretation of evolutionary responses (Kawecki *et al*., 2012), it seems unlikely to us that this effect would have strongly contributed to the differences observed here in only three generations, not least because the same census population sizes were maintained across time and populations, and because the differences in the mating patterns did not involve strong inbreeding, which would have had a greater impact on the effective size.

In summary, our experiment revealed rapid evolutionary changes in vegetative growth following evolution of a wind-pollinated plant under contrasting densities, which occurred in only three generations. The results of our experiment join those of several others that demonstrate how responsive to selection experimental plant populations can be. These studies have focused on a number of traits related to plant reproduction, e.g., floral scent production, the ability to self-fertilize (Gervasi & Schiestl, 2017; Schiestl & Johnson, 2013; Ramos & Schiestl, 2019), sex allocation (Dorken & Pannell, 2009; Cossard *et al*., 2019), and pollen performance abilities (Lankinen *et al*., 2017). Our study now shows how a modification of growth conditions during the reproductive period may result in rapid changes to vegetative traits, too (and see Schiestl & Johnson, 2013). Perhaps the most striking aspect of our findings is the demonstration of evolutionary responses in traits expressed early in a plant’s life as a result of differences in reproductive success expressed at the time of reproduction, when seeds were sired by males and produced by females. As we have explained, it seems likely to us that responses by males and females were mediated by selection via competition for mates and for light, respectively. Finally, whether or not this interpretation is accurate, our results overall point to a complex interplay of different modes of selection that are evidently sensitive to a key demographic variable, plant density. Future studies might resolve some of this complexity through experiments that vary sex ratios or the number of pollen donors more directly that we have done here.

## Acknowledgments

We thank Guillaume Cossard, Jeremy Devaux, David Degueldre, Pauline Durbin, Jacqueline Llorca, Estelle Barbot, Guillaume Rubi, Augustin Chen and Célia Morillas for their technical assistance and the team of the “Plateforme des Terrains d’Expériences du LabEx CeMEB” (Montpellier, France), which hosted part of the experiment. We thank Céline Teplitsky for helpful advice on the comparisons of G matrices. JT was supported by a grant to JRP by the Swiss National Science Foundation (31003A_163384) and by a by a Marie Skłodowska-Curie grant (#844321) to JT.

---

**Figure S1**. Predicted sex-specific branch length in evolved and source populations of *Mercurialis annua* grown in a common garden after three generations of evolution. Branch length was treated as a response variable in our null models, which included both block and sex by population random effects. Females and males are represented by pink triangles and blue circles, respectively. The significance of differences between treatments (source, low-density and high-density) in models combining both sexes was evaluated using LRTs (\**p* < 0.05). Horizontal bars indicate standard errors in model estimates.

**Figure S2:** Phenotypic correlation matrices with Pearson correlation coefficients and their associated p-values between pairs of vegetative and reproductive traits. Phenotypic matrices were calculated by pooling individuals from the source populations (males: a. and females: d.), from the low-density treatments (males: b. and females e.) and from the high-density treatment (males: c. and females: f.). Measured vegetative traits were common between males and females and are displayed in red for the purpose of inter-sex comparison. Pearson correlation coefficients are illustrated using color shading, with blue and red corresponding to positive and negative coefficients, respectively (as shown by horizontal legend bars). Significance of each correlation coefficient was evaluated (. p<0.10, ^*^p <0.05, ^**^p<0.01, ^***^p<0.001).

**Figure S3**. Mean positions along the first eigentensor and their credibility intervals extracted from a tensor analysis comparing the three phenotypic correlation matrices of the three different treatments for males (a) and females (b). Credibility intervals were estimated on the basis of 1,000 phenotypic correlation matrices calculated by re-sampling measured individuals. Separate for each sex, the first eigenvector accounted for respectively 91% and 99% of the variation between the three matrices in males and in females.

## Notes

### Competing Interest Statement

The authors have declared no competing interest.

