## Supplementary files for "Rapid divergence in vegetative morphology of a wind-pollinated plant between populations at contrasting densities"

**Table S1** : Testing for (a) spatial structure in plant reproductive traits and (b) evolutionary response in these reproductive traits of *Mercurialis annua* plants that evolved at high- *versus* low-density and compared to our source population (SP) over the course of three generations, as assessed in a common garden.

| | Plant trait | (a) Spatial structure | | (b) Treatment effect | | | | | | | |
|---|---|---|---|---|---|---|---|---|---|---|---|
| | | (df=3) | | (1) SP – High – Low (df=2) | | (2) SP – High (df=1) | | (3) SP – Low (df=1) | | (4) Low – High (df=1) | |
| | | $\chi^2$ | p | $\chi^2$ | p | $\chi^2$ | p | $\chi^2$ | p | $\chi^2$ | p |
| Males | Peduncle length | 5.88 | 0.12 | 0.62 | 0.73 | 0.0218 | 0.88 | 0.0718 | 0.79 | 0.62 | 0.43 |
| Males | Number of peduncles | 3.41 | 0.33 | 0.496 | 0.78 | 0.330 | 0.57 | 0.360 | 0.55 | 0.0512 | 0.82 |
| Males | Number of peduncles on top | 5.85 | 0.12 | 3.20 | 0.20 | 3.38 | 0.066 | 1.33 | 0.25 | 0.603 | 0.44 |
| Males | Total peduncle mass | 7.77 | 0.051 | 1.64 | 0.44 | 1.34 | 0.25 | 1.80 | 0.18 | 0.0059 | 0.94 |
| Females | Seed number | **11.6** | **0.00889** | 0.58 | 0.75 | 0.176 | 0.68 | 0.542 | 0.46 | 0.200 | 0.65 |
| Females | Seed size | $3.33 \times 10^{-9}$ | 1.00 | 0.893 | 0.64 | 0.475 | 0.49 | 0.934 | 0.33 | 0.107 | 0.74 |
| Females | Total seed mass | **8.43** | **0.038** | 0.723 | 0.70 | 0.174 | 0.68 | $9.00 \times 10^{-4}$ | 0.98 | 0.646 | 0.42 |

**Notes:** The spatial structure for plant traits was evaluated by constructing models that explained them as a function of a spatial random effect modeled by a Matérn function, including three parameters. The effect of the treatment was assessed by comparing models thanks to LRTs that included or not the treatment effect which incuded : (1) source population, experimental populations evolved at low and at high density, (2), source and high-density populations, (3) source and low-density populations and (4) low- and high-density populations. Models were fitted by maximum likelihood for performing LRTs between models differing in their fixed-effects structure, and by restricted maximum likelihood for LRTs between models differing in their random-effect structures.

Significant p-values are highlighted in bold and degrees of freedom (df) are provided for each type of LRT.

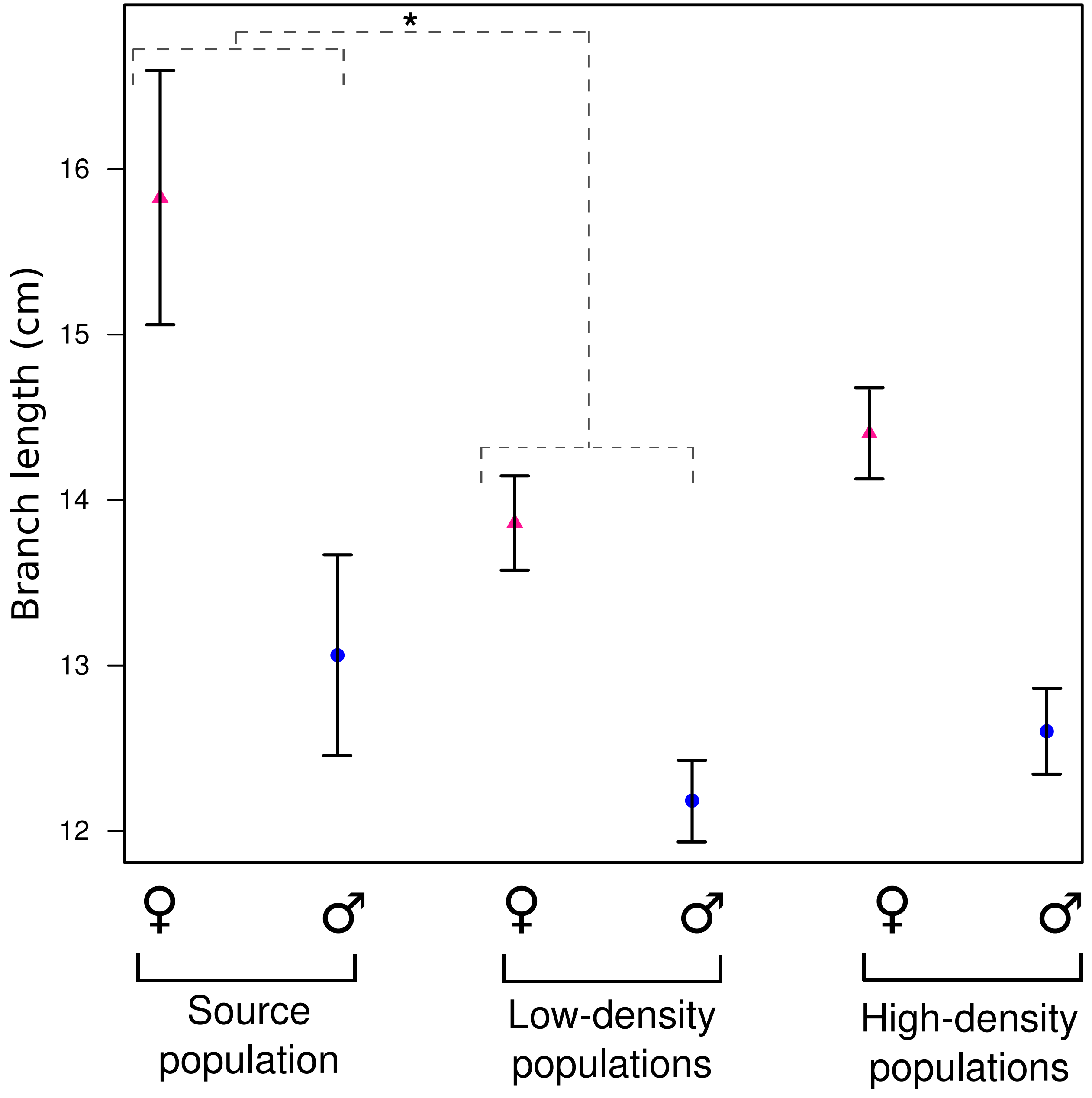

*
16
15
14
13
12
Branch length (cm)
♀
♂
♀
♂
♀
♂
Source population
Low-density populations
High-density populations


Figure S1

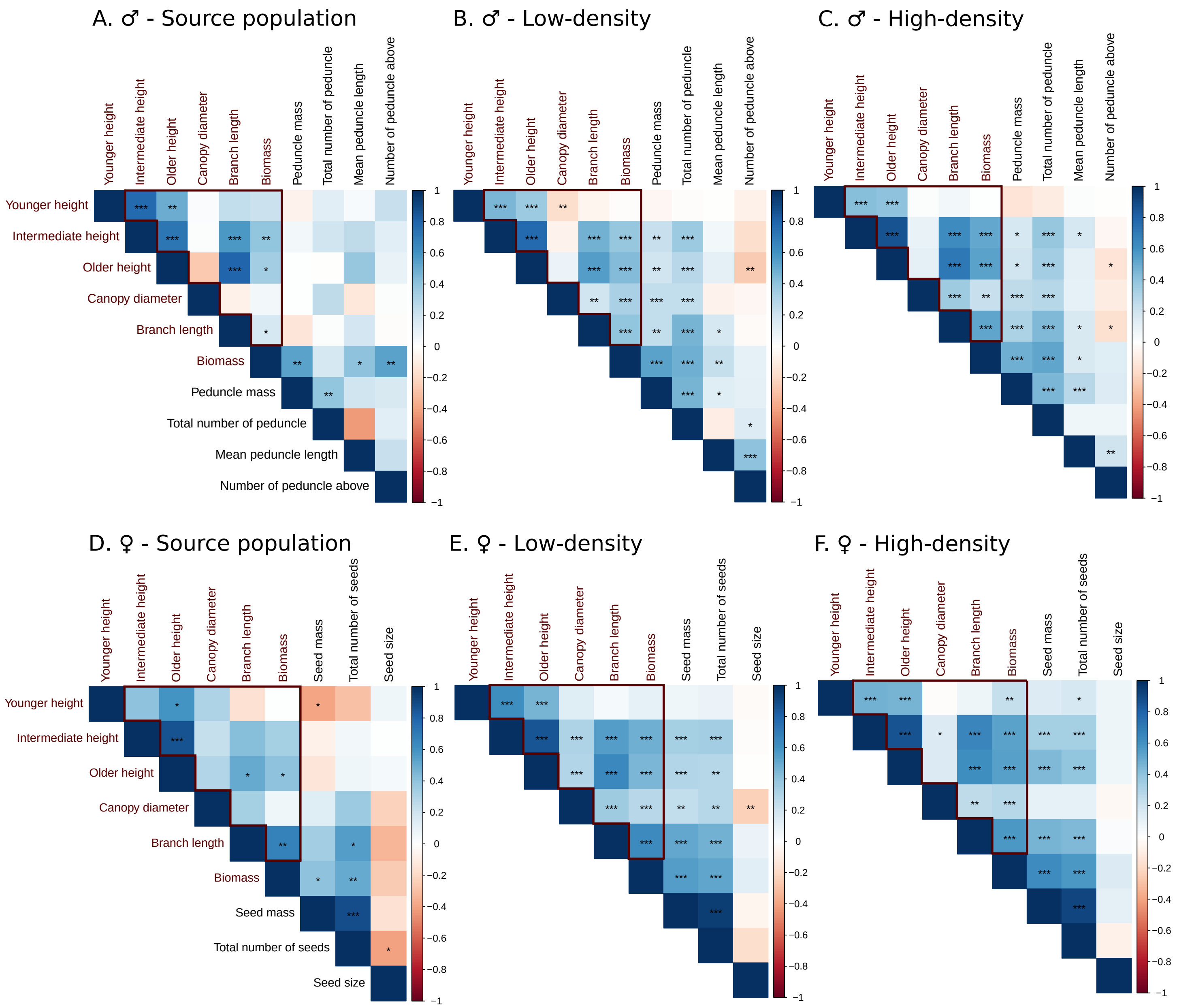
A. ♂ - Source population
B. ♂ - Low-density
C. ♂ - High-density
D. ♀ - Source population
E. ♀ - Low-density
F. ♀ - High-density
Younger height
Intermediate height
Older height
Canopy diameter
Branch length
Biomass
Peduncle mass
Total number of peduncle
Mean peduncle length
Number of peduncle above
Seed mass
Total number of seeds
Seed size

Figure S2

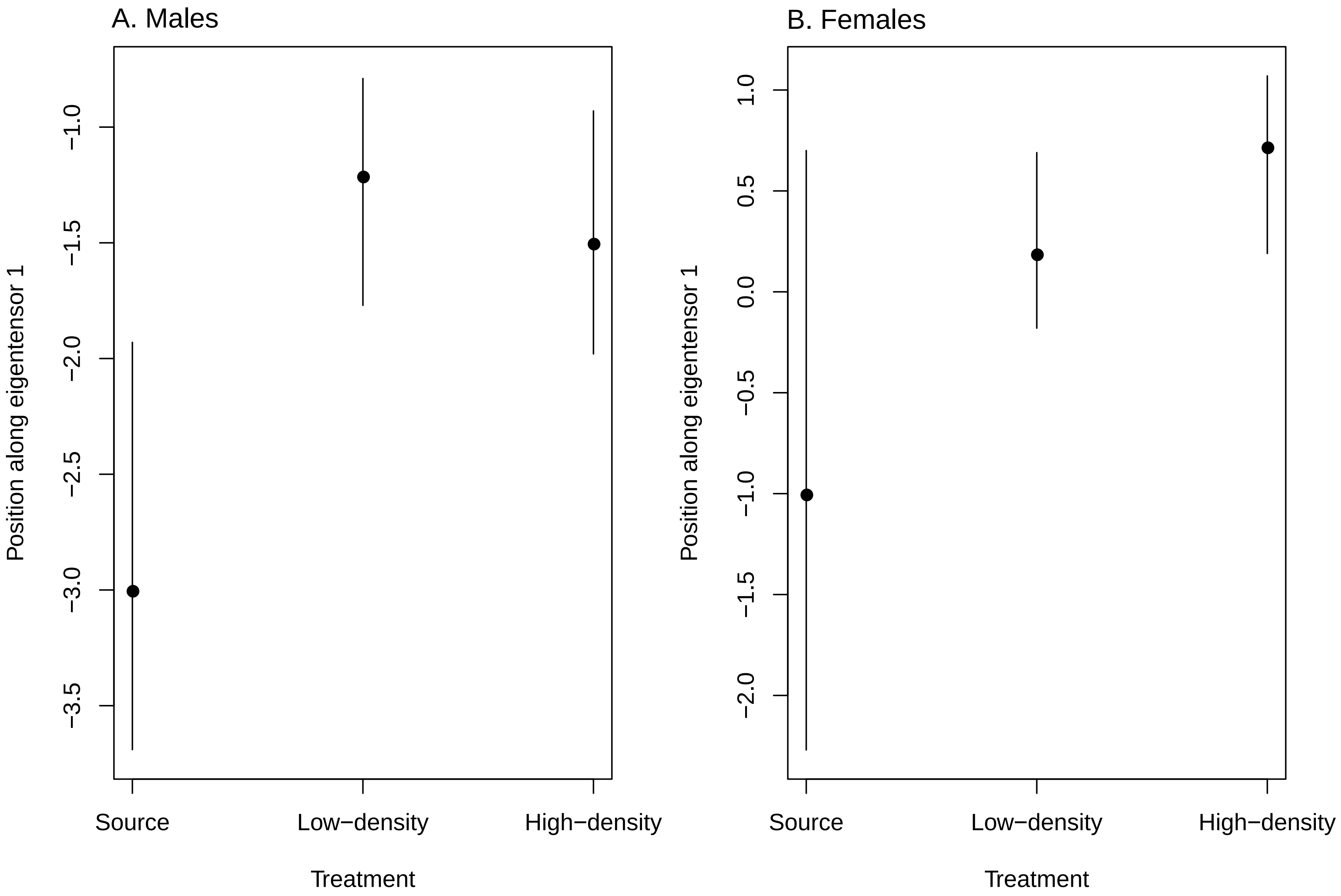
A. Males
B. Females
Position along eigentensor 1
−1.0
−1.5
−2.0
−2.5
−3.0
−3.5
1.0
0.5
0.0
−0.5
−1.0
−1.5
−2.0
Source
Low−density
High−density
Treatment

Figure S3
